# Flight height patterns of a critically endangered insectivorous bat, impacted by wind turbine collision

**DOI:** 10.1101/2025.05.23.655017

**Authors:** Amanda Bush, Lindy Lumsden, Thomas A. A. Prowse

## Abstract

**Background:** Renewable energy production is being developed worldwide to reduce reliance on fossil fuels and thereby moderate the rate of anthropogenic climate change. Harnessing wind energy, using wind turbines, is a prominent form of renewable energy production. There are, however, biodiversity impacts, including collisions by birds and bats with rotating blades. High levels of mortalities can cause localised and species-level population declines, which is especially significant for threatened species. The height at which species fly is a key collision risk factor. In this study we investigated the flight height patterns of a critically endangered bat to inform mitigations to reduce impacts.

**Methods:** We captured and GPS tagged 244 Southern Bent-wing Bats (*Miniopterus orianae bassanii*) (14 – 19 g) in south-eastern Australia, in spring and late summer/early autumn. We retrieved 93 units, yielding 4,289 bat observations from an 18–21 day period in each season. The vertical measurement error of the GPS units at different heights and sampling intervals was investigated with drone test flights. A Bayesian state-space modelling approach was then developed to estimate flight height distributions while accounting for measurement error.

**Results:** Our results suggest that the majority of bat activity occurs between ground level and 30 m altitude, at least in early spring and late summer/early autumn. However, the bats were recorded taking short flights above 60 m and at times even flew above 80 m (maximum model estimate was 92.7 m with a maximum 95% CI of 144.1 m) and demonstrated that they are capable of quick and frequent altitudinal changes from ground level to almost 40 m. Flight heights were on average higher when associated with trees in summer. Mean flight heights appear to be higher when associated with treed habitats in summer.

**Conclusions:** This study shows that the flight height distribution of small bats can be investigated using store-on-board GPS devices and illustrates a statistical approach that incorporates vertical measurement error. It provides the first insights into the flight heights of this small bat species, which can help inform a more complete flight height profile across different seasons and conditions with future improvements in GPS technology. Results suggest that, although the Southern Bent-wing Bat primarily flies at lower heights, it exceeds 30 m altitude at times, increasing the risk of mortalities due to wind turbine collision. Accurately determining the proportion of time that threatened species such as the Southern Bent-wing Bats spend within high-risk flight zones will provide the evidence base for implementing effective mitigation measures to reduce population-level impacts.

## BACKGROUND

There is an increasing focus worldwide on renewable energy production as a means of reducing reliance on fossil fuels^1^. A prominent form of renewable energy comes from harnessing wind energy through the use of wind turbines. However, there is growing concern regarding the negative consequences for volant (i.e. flying) birds and bats that die or are injured after colliding with rotating wind turbine blades ^2–4^. In Europe and North America, there is evidence that the cumulative effects of individuals colliding with wind turbines may impact species viability at a population level ^5,6^. Local-scale population impacts are also a concern ^7,8^. A key issue is the extent to which individuals are vulnerable to wind turbine collisions based on the height at which they fly.

Bird and bat species fly at varying heights above the land surface, which is influenced by foraging strategies, size and wind morphology ^9,10^. Flight height and time spent at particular heights also vary within a species due to the purpose of the flight (e.g. local commuting, migrating or foraging/ hunting, weather conditions, topography, habitat and season) ^11–15^. These factors in turn influence a species’ or individual’s risk of collision with rotating turbine blades ^15–17^. Understanding the flight height patterns of birds and bats can inform an assessment of turbine collision risk. This can then enable proactive measures like turbine curtailment during periods of high collision risk to be implemented to better manage this risk.

The flight heights of insectivorous bats have typically been investigated by recording their ultrasonic calls at different heights, by fastening acoustic recorders to structures or anchored balloons ^16,18,19^. Although these methods can be informative, the area of airspace that can be sampled is restricted due to the height and number of available structures or balloons, the short reception range of ultrasonic recorders, weather-related sound attenuation, and the species-specific detectability distances of bats with different call characteristics ^20^. Furthermore, calls may not be readily identifiable as some bats change their call structure at higher altitudes ^21,22^, or there may be interference from wind noise ^20^. These limitations can make it difficult to accurately quantify the relative activity of different species at different heights, and also make it difficult to assess flight heights across the spectrum of weather conditions over which bats may be active. Radar is another option for investigating flight heights of insectivorous bats ^23^ but bats as a group are difficult to distinguish from birds and insects ^24^, and also individual species cannot be identified from radar images alone. Flight heights of insectivorous bats vary depending on the species and specific behaviours, but bats have been recorded from just above ground level to more than 3,000 m in height ^23,25,26^.

Increasingly flight heights are estimated by using barometers, altimeters, geolocators and GPS trackers fitted to target species ^27–31^. GPS units in particular enable recorded heights to be associated with different habitats (as GPS also records location in a horizontal plane), providing a more nuanced picture of species habitat use. GPS units estimate locations by triangulating information received from multiple satellites ^32^, with more battery-efficient sampling methods providing a “snapshot” of satellite information for post-processing after data download ^33^. The positional accuracy of GPS locations is influenced by the number of satellites detected and the position of those satellites relative to each other ^34^, with closer grouping of satellites yielding a greater degree of location error^35^. GPS units are more accurate in situations where they have a clear view of the sky, with denser forest canopy reducing accuracy in both the horizontal ^32^ and vertical planes ^36^. These factors potentially reduce location and height accuracy of individuals that fly below or within the tree canopy. Other factors that influence accuracy include flight speed (stationary versus moving), unit sampling frequency ^28^ and even the use of different units of the same GPS model ^34^. The precision errors of GPS fix locations are often only given in the horizontal plane with the Horizontal Dilution of Precision (HDOP), indicating the precision of the achieved latitude and longitudes, with some units also recording a Vertical Dilution of Precision (VDOP). GPSs are inherently less accurate in the vertical plane ^37^ but flight heights are still at times reported verbatim in the literature without considering this additional source of error (e.g. ^38^). At times implausible values (e.g. negative heights) are adjusted ^26^ or removed, potentially introducing a positive bias to resulting mean flight heights, and losing information about measurement error in the process ^34,37^ .

Bayesian state-space models that partition process error (i.e., true variation in flight heights or ‘states’) from measurement error are useful for modelling flight height data obtained from GPS tracking studies. They provide a robust method to simultaneously consider the mean and variance of flight heights, which is critical for estimating the proportion of time bats spend within different height zones. These models specify a probability distribution that generates measurement errors, and prior knowledge of this distribution can assist with disentangling the process and error components. Previous GPS-based studies have assumed measurement errors arise from a normal or t-distribution, potentially with parameters (e.g. the variance) governed by DOP. ^27,34,37^

Bayesian state-space models have been adopted for modelling flight height from GPS tracking studies of large bird species (including raptors and sea birds), typically using many thousands of GPS locations ^27,34,37^. However, it is much harder to estimate flight height distributions for small insectivorous bats (<20 g) because they can only carry lightweight store-on-board GPS units that must be retrieved to obtain the data. The small size of commercially available units and batteries limits the volume of three-dimensional data that can be collected from a single individual to at most several hundred fixes ^39,40^, making long-term, multi-season data collection difficult, expensive and resource intense. These limitations necessitate a sampling regime that balances sampling interval with length of operation. The lower GPS fix frequencies used to achieve this balance may result in higher levels of vertical measurement error ^28^ which warrants careful consideration when assessing the flight heights recorded for free-flying bats. Perhaps because of these additional challenges, the field of estimating flight height distributions for small insectivorous bats from GPS tracking data is in its infancy.

The Southern Bent-wing Bat (*Miniopterus orianae bassanii*, ‘SBWB’ herein) is a Critically Endangered obligate cave-roosting insectivorous bat that is endemic to south-eastern Australia. It breeds in only three maternity roosts, in the states of Victoria and South Australia each spring/summer ^41^, but moves regularly between maternity and non-breeding roosts year-round ^42^. Much of the SBWB’s distribution overlaps with high-value wind resources in southern Australia ^43,44^. SBWB mortalities have been documented from multiple wind energy facilities, indicating that the species flies within the rotor swept area of operational turbines at times ^45,46^. The flight height profile of the species is however poorly understood.

Here, we use miniaturised GPS technology to investigate the flight height patterns of SBWB (hereafter ‘bats’ for bats fitted with GPS trackers) over different habitats and seasons in a varied landscape. Flight heights were compiled from GPS tracking studies of three subpopulations in Victoria and South Australia (based around the maternity sites near Warrnambool, Portland and Naracoorte) and referenced to the habitat type at the GPS fixes. We also flight tested GPS units on a multi-rotor drone using actual and modelled bat flight paths, and used a Bayesian state-space modelling approach to investigate the error associated with vertical height measurements. This approach overcomes some of the uncertainties associated with using lightweight GPS trackers (<1.5 g) to estimate the flight height patterns of small insectivorous bats, and may be more widely applicable to other small bat species, especially with future improvements in this technology.

## METHODS

### GPS tracking of SBWBs from multiple subpopulations

We conducted GPS tracking of individuals from the three subpopulations of the SBWB in south-eastern Australia. The landscape in the study region is variable, ranging from open agricultural lands with scattered trees and planted windrows to plantation and native forest. Tracking occurred across five different time periods: in February/March 2020 and 2021 for the Warrnambool subpopulation, in February/March 2023 for the Naracoorte subpopulation, and in September 2023 and February/March 2024 for the Portland subpopulation. Collectively the tracking occurred across four separate years and two seasons, and across 42 days of the year (21 in spring and 21 in summer-autumn), with the majority of data collected in late summer and early autumn (hereafter ‘summer’), after the breeding season.

During each deployment, adult bats of both sexes were fitted with paired GPS/VHF telemetry units. The GPS components were Lotek (Wareham UK) Pinpoint 10 units (1.0 g) which store all data on-board. The VHF transmitters used were Lotek PicoPip Ag337 (0.29 – 0.32 g) or Holohil (Carp, Canada) BD-2X transmitters (0.35 g). The VHF transmitters assisted with locating the GPS units to enable the data to be retrieved once they had fallen off the bats. Small pieces of reflective tape were added to the units to assist with finding them in cave roosts. We glued the paired GPS/VHF units to captured bats using a non-toxic adhesive designed for human use (Urobond IV or V, Urocare Inc, USA). The combined weight of units, tape and glue was ≤ 1.5 g, which was < 10 % of the body weight of those bats selected for the study. At some sites a purpose-designed automated VHF receiving system was used to detect tagged bats and provide daily reports via email to assist with unit retrieval.

The Lotek GPS units were programmed to take SWIFT fixes, which involves the unit taking a snapshot of satellite locations, and using post-processing algorithms to calculate 3-dimensional location data^32^. Fix attempts timed out if a location could not be obtained within 12 seconds. We varied the scheduled frequency of fixes to obtain movement information at different resolutions. Samples were variously taken at 1-minute (34.2 % of all fixes), 2-minute (9 % of all fixes), 15-minute (2.2 % of all fixes), and 1-hour intervals (54.6 % of all fixes). Depending on the scheduled sampling interval and battery performance, units sampled for *c*. 2-4 hours in a single night, or for 8-9 hours on up to 18 nights. The GPS units recorded the latitude and longitude (WGS 84 ellipsoid) of each individual during each successful fix, as well as height above sea level and the number of satellites used to estimate the horizontal and vertical position. They also provided two measures of accuracy: Horizontal Dilution of Precision (HDOP) and ‘eRes’ (Lotek’s proprietary measure of the accuracy for SWIFT fixes). We fitted 244 units and retrieved usable data from a total of 93 bats: 22 bats from the Warrnambool subpopulation (12 males and 10 females), 44 bats from the Portland subpopulation (23 males and 21 females) and 27 bats from the Naracoorte subpopulation (12 males and 15 females).

### Data processing and environmental covariates

All data processing and analyses were carried out in R vers. 4.2.2 ^47^. GPS locations with a HDOP measure greater than 6 were excluded from subsequent analyses to minimise horizontal error that could compromise the classification of habitats beneath these locations. Data for individual bats with fewer than six valid GPS locations were also excluded. This resulted in 95.7 % of the original dataset (4298 fixes) being retained and used for analysis.

The height above land (hereafter ‘raw flight height’) for each position fix was calculated by subtracting the ground elevation sourced from a digital elevation model (DEM) from the GPS tracker recorded altitude for each bat location. The ground elevation was extracted based on the latitude and longitude using the ‘raster’ package ^48^. Ground elevation was extracted from separate DEMs for Victoria (vertical error = +/- 5 m [root mean square error], horizontal error = +/- 12.5 m, 10 m resolution) and South Australia (5 m resolution, vertical and horizontal error unknown). Where location fixes were sited over the ocean, the underlying elevation was set to zero to represent sea level, as these points were outside the terrestrial DEMs. Each fix was categorised as being either associated with trees (treed) or not associated with trees (untreed) by extracting point location vegetation data from the Digital Earth Australia Land Cover (30 m resolution) layer ^49^. The treed classification includes Natural Terrestrial Vegetated Woody (closed or open) or Natural Aquatic Vegetated Woody. Woody vegetation has at least 20 % canopy cover, and therefore the dominant vegetation is likely to consist of both shrubs and trees. There is no information available regarding the height of the vegetation. All other categories including Natural Terrestrial Vegetated Herbaceous, Cultivated Terrestrial Vegetation and water were categorised as untreed. The sunset time for each night of data collection was calculated at an average location of maternity cave roosts using the ‘suncalc’ package ^50^.

### GPS drone testing to estimate vertical error

To assess the vertical positional error of the GPS units, we conducted a three-day validation trial in late July and early August 2024, using a subset (n = 10) of the GPS units from the study to fly on a drone. We used a DJI Matrice 350 RTK drone connected with a DJI D-RTK 2 base station to minimise positional error (both vertical and horizontal) of location data. To ensure high accuracy of the drone position during flights, the location of the base station was surveyed with an Emlid Reach RS3 RTK-GPS to obtain an accurate (approx. 1-2 cm error) absolute position. The base station provided a precise reference for the drone’s position with the expectation of a positional error of approximately 2 cm horizontally and 4 cm vertically ^51,52^. The drone typically connected to more than 30 satellites during the trial flights. The drone recorded positional data in the WGS 84 ellipsoid projection. Four GPS trackers were fixed (approximately 10 cm apart) to a custom-built carbon plate positioned 20 cm above the body of the drone to minimise interference between the drone and the trackers. The drone was operated by a pilot from The University of Adelaide’s Uncrewed Research Aircraft Facility.

We used both the actual bat flight data from the GPS tracking (one-minute sampling frequency, ‘bat flights’ herein), as well as several simulated flightpaths (‘modelled flights’) to provide flight speeds, altitudes and turning angles for the drone flights. To inform the drone’s ‘bat flight’ patterns, we selected six 20-minute GPS series (horizontal coordinates and raw heights) obtained from free-flying bats that represented a mix of relatively direct flights (where bats were continually moving to new locations) and instances where bats were making more localised movements in a discrete area.

These were combined into three 40-minute flight plans. In contrast, the ‘modelled flights’ were simulated using a discrete-time autoregressive moving average (ARMA) process for flight height and speed across the ground. We assumed a mean flight height and speed of 20 m and 4 m/s, respectively, and set the ARMA parameters to produce a distribution of flight heights and speeds that was similar those observed in tracked bats. We also assumed a time step between successive locations of 10 seconds and randomly sampled horizontal turning angles between locations from a normal distribution with mean zero and standard deviation of 30 degrees.

Where necessary, we made the following adjustments to the GPS series used to program the drone. Firstly, the horizontal coordinates were adjusted to fit within one of two *c*. 500 m^2^ areas of relatively open field within The University of Adelaide’s Roseworthy Campus. Due to safety reasons, each area was in open farmland with a clear view of the sky, with no nearby buildings and just a few low sparse trees. To adjust the data series, the starting coordinate of each flight was rescaled to the centroid of the paddock polygon, and then the observed turning angles and ground speeds from the GPS series were used to generate a series of 3-dimensional locations. However, where ‘bat flights’ would have moved beyond the testing area, the next position was calculated by reflection from the area boundary using a random turning angle, whilst maintaining the true distance travelled. Secondly, given aviation regulations prohibiting drone flights above 120 m and collision risks associated with very low trajectories, we ensured the drone remained between 10 and 120 m above land by truncating the observed raw flight heights values falling outside this range (to the lower or upper limit as necessary). Where bat flight speeds were below the minimum flight speed capability of the drone (3.6 km/h) they were adjusted to this speed.

The testing occurred between approximately 12:00 and 16:15 on day 1, 10:30 and 16:15 on day 2, and 10:00 and 15:30 on day 3. Test flights lasted up to 40 minutes depending on the rate of drone battery drain, which was influenced by wind conditions. We varied the GPS sampling interval (1-minute, 2-minute, 7-minute or 15-minute intervals) to investigate if sampling frequency impacted vertical measurement error ^28^, at least over these shorter sampling intervals. However, it was impractical to consider vertical measurement error over the longer sampling intervals (up to 1 hour) that were used when tracking the bats. Each GPS unit was tested at each frequency multiple times throughout the trial. The drone remote controller reported the drone’s position nine to 11 times each second based on its position relative to the base station. Each bat flight scenario and modelled flight were flown on between two and six occasions over the three-day period, and at different times throughout the day to account for changing positions of the orbiting satellites.

### Data Analysis

#### Drone trial

Estimating the vertical error associated with the bat GPS units required precise matching of the drone and GPS vertical positions over the three-day trial period. We initially conducted this matching by identifying the closest pairs of timestamps from each device (in milliseconds). However, we also tested different time lags between the GPS measurement and drone positions and evaluated the horizontal error obtained for each lag (due to the lower error associated with latitudinal and longitudinal data than altitude from GPS units). This approach identified a 2.5 second time discrepancy between the GPS measurement and drone position, which produced the lowest median horizontal error across the different time shifts tested. This likely resulted from the time taken for the GPS units to initiate and complete the process of recording a position fix. Therefore, we calculated vertical measurement error for each GPS location as the GPS height measurement minus the drone height from 2.5 seconds prior.

We investigated the bias and variance of the vertical measurement error process through a bespoke Bayesian model assuming these errors *v*_*t*_ arose from a Student t-distribution

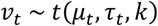

where *k* was set equal to 4 to ensure finite mean and variance. With *k* set thus, μ_*t*_ is the time-varying mean vertical error that represents the bias, and the precision τ_*t*_ is simply the inverse of the variance σ_*t*_^2^. We initially developed separate linear models for the bias and standard deviation

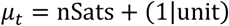

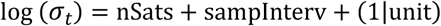

where *nSats* is the number of satellites contacted for the GPS location fix, *sampInterv* is a categorical variable representing the different sampling intervals programmed, and 1|unit represents the random effect of each individual GPS unit tested. However, given no strong evidence of a difference in standard deviation due to different sampling intervals (Figure S1), we excluded this term and refitted the model for the precision.

We developed the model using the BUGS language, which was implemented using Bayesian Markov chain Monte Carlo in JAGS version 4.3 ^53^, and which we ran through the functions in the R package ‘rjags’ ^54^. We ran three Markov chains for 5,000 iterations each, with a burn-in of 2,000 iterations and a thinning rate of 5 to reduce autocorrelation in the posterior samples. Model convergence was assessed through visual inspection of chains and the Gelman-Rubin statistic (values close to one indicate convergence) ^55^.

#### Flight height distribution model

We developed an integrated Bayesian state-space model to analysis the distribution of flight heights of bats above the ground surface. The goal of our state-space model for observed flight heights above land (*y*) was to partition variation in the true heights (ℎ) from variation introduced by the imperfect measurement process ^56^, while also investigating drivers of variation in the mean flight height. The model integrated data from the drone testing to inform estimation of the measurement error process for the tracked bats. We developed the Bayesian model using the BUGS language (Supplementary Appendix A1) for implementation in JAGS software ^53^.

The first component of our state-space model is the process model

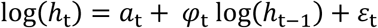

which represents a log-scale first-order autoregressive (AR1) process linking the current flight height (ℎ_*t*_) to the previous height (ℎ_t-1_) through the correlation parameter φ_*t*_, and where ε_*t*_ is the normally distributed process error term. Given we used different GPS sampling frequencies when tracking the bats, we assumed a conditional autoregressive (CAR) process governing how the strength of φ decayed exponentially as a function of the time interval (ω_*t*_, minutes) between successive fixes

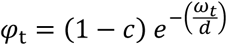

where *c* and *d* are estimated parameters.

The intercept term *a_*t*_* in the first equation above is equal to *y*-_*t*_(1 − φ) where *y*-_*t*_ is a time-varying mean flight height, thereby allowing a model for *y*-_*t*_ through manipulation of *a*_*t*_. We expected bat flight heights might be influenced by habitat (treed or untreed vegetation), season and the sex of the tracked individual, along with the time since sunset. We therefore modelled variation in the mean flight height as

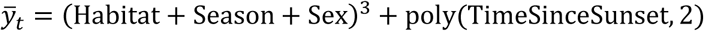

where the cubic power represents all main effects, including interaction terms for the three categorical variables, and the final term represents a 2-order orthogonal polynomial allowing a potentially non-linear effect of time since sunset. Mean flight heights were generated for all categories at the mean time since sunset (245 minutes after sunset each night).

The second component represents the GPS measurement error process, which is linked to estimates of the true flight height (ℎ_*t*_) but is affected by the stochastic error term (*v*_*t*_)

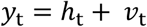

This equation assumes there is no consistent bias in the altitude measurements obtained from the Lotek GPS units. As detailed above, we assumed the errors (*v*_*t*_) arose from a Student’s *t*-distribution with a mean of zero, and precision (τ) and degrees of freedom (*k*), which were estimated from the data. We also assumed a non-linear (quadratic polynomial) relationship between the parameters τ and *k* and the number of satellites used for a GPS location fix, which integrated data from both the drone tests and the bat datasets.

We fitted the state-space model using Bayesian Markov chain Monte Carlo in JAGS version 4.3 ^53^ which we ran through the functions in the R package ‘rjags’ ^54^. Uninformative Normal (0,1000) priors were used for fixed effects, while Uniform (0,10) priors were applied to random-effects variances for model identifiability. We ran three Markov chains for 50,000 iterations each, with a burn-in of 10,000 iterations and a thinning rate of 5 to reduce autocorrelation in the posterior samples. Model convergence was assessed through visual inspection of chains and the Gelman-Rubin statistic (values close to one indicate convergence) ^55^.

## RESULTS

### GPS Data

A total of 4,289 fix locations were collected from bats, from between 19 – 21 days each, and across each of the four summer periods (between February 5^th^ and March 5^th^) and for 21 days in early in spring (September 9^th^ to 29^th^). These consisted of 236 location points for males and 204 for females in spring, and 1821 location points for males and 2028 for females in summer. There were fewer data points later in the sampling periods as units either detached from the bats or ceased sampling. Raw flight heights above estimated ground elevation (GPS altitude output minus elevation from the relevant DEM) ranged from -135 m to 565 m, and 13% of the points were recorded as being at ≤0 m. The range of ground elevations that bats flew over was between sea level and 240 m.

### Drone trials

The drone validation flights illustrated substantial vertical measurement error associated with the Lotek GPS units used for our study. The GPS-measured flight heights ranged from 19.4 m to 269.7 m above sea level, whereas the true drone heights only ranged from 75.6 m to 192.7 m. The mean vertical error was 10.2 m, and 81.8 % of observations fell within 20 m of the true vertical position. Preliminary modelling of the bias and standard deviation of the t-distributed vertical measurement error process demonstrated a strong influence of the number of satellites contacted, but no impact of sampling interval. Therefore, we focus our reporting on a model without inclusion of the latter term. There was a positive bias observed in the GPS height measurements of around 10 m (Figure 1a,c) which was negatively but not significantly related to the number of satellites contacted (coefficient estimate [± 95% credible intervals] = 2.81 [-5.82, 0.16]). In contrast, there was a strongly negative and significant relationship between the standard deviation of this error process and the number of satellites contacted (log-scale estimate: -1.56 [-1.82, -1.30]) (Fig 1b,c).

**Figure 1.**
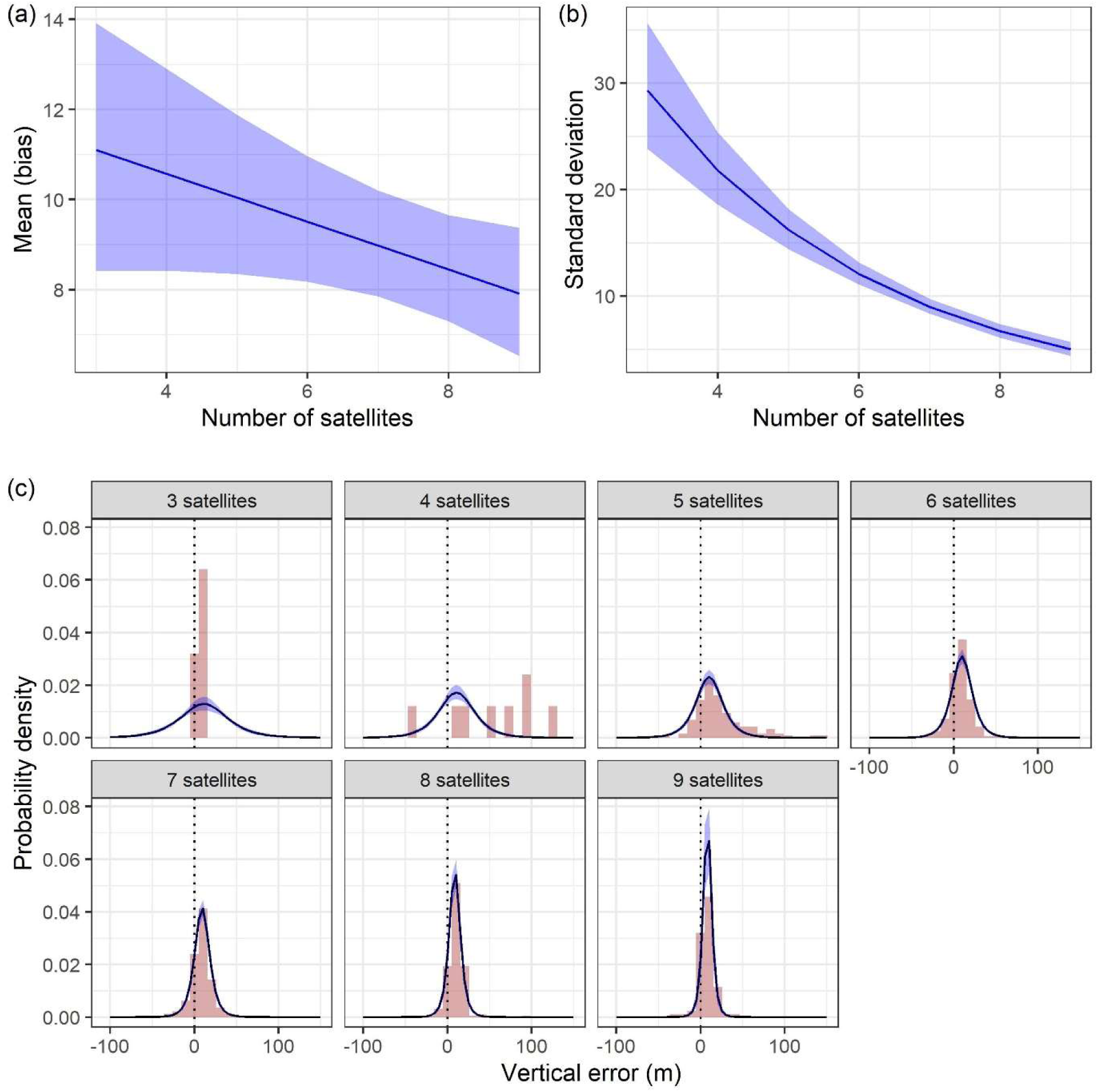
Drone trials. (a) Mean (bias) and (b) standard deviation of the GPS vertical measurement error process as a function of the number of satellites contacted for a location fix. Blue shading represents the 95% credible intervals for these relationships. (c) The estimated t-distributions (black solid lines) capturing vertical measurement error for different numbers of satellites contacted. The background histograms (brown shading) illustrate the distribution of observed vertical errors, while the dotted vertical line indicates zero vertical error.

### Flight height patterns from raw and modelled data

The distribution of flight heights for both the raw and modelled heights suggest that the majority of SBWB activity is occuring between ground level and 30 m, during our sampling in summer and early spring (Table 1). It also indicates that there is a general pattern of decreasing flight activity with increasing height above ground level.

**Table 1.** Percentage of raw and modelled flight height points from all data that are above the specified height above estimated ground level.

|  | Raw flight heights | Modelled flight heights with 95% credible intervals |
| --- | --- | --- |
| -30 m | 99.370 | - |
| 0 m | 86.990 | 100 |
| 30 m | 8.697 | 4.024 [2.751, 5.642] |
| 60 m | 2.192 | 0.262 [0.104, 0.489] |
| 90 m | 0.886 | 0.043 [0, 0.093] |

### Mean flight heights

In summer the mean modelled flight height (at the mean time since sunset - 245 minutes after sunset) was greater above trees that in untreed habitat (Figure 2a) and in spring the mean flight height of males in untreed habitat was lower than all other categories. The mean modelled flight heights ranged between 1.2 m (males, untreed in spring) and 13.7 m (males, treed in summer) across all the categories, with the highest 95 % credible interval of the mean reaching 15.2 m (males, treed in summer). The modelled mean flight height across the night for all seasons, sexes and habitat classifications combined was relatively consistent (Figure 2b).

**Figure 2.**
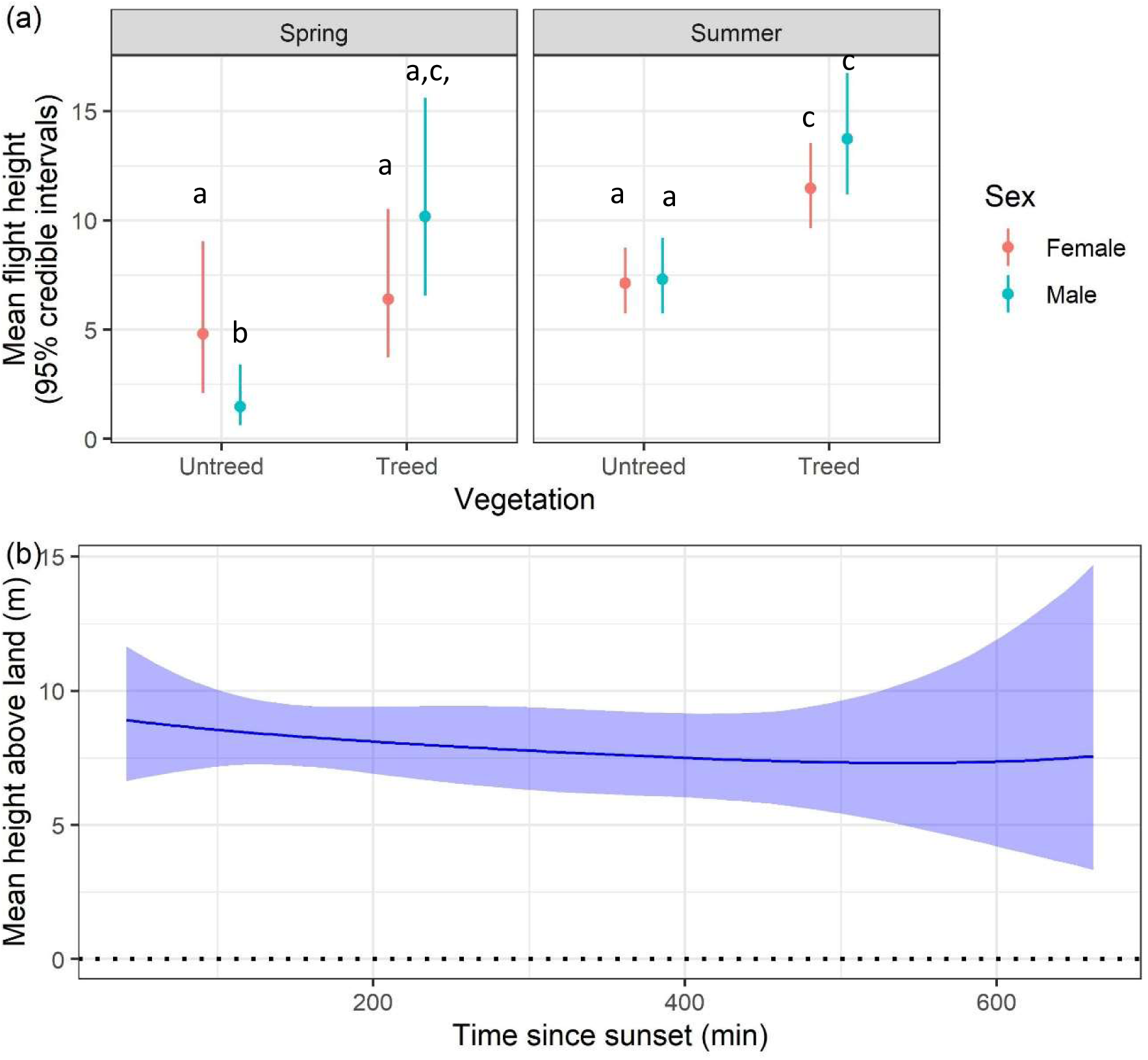
(a) Modelled mean flight heights and 95% credible intervals in meters above estimated ground level by sex and season at 245 minutes after sunset (mean time since sunset across all nights). Habitat at each point location is classified as either treed or untreed based on a 30 m resolution tree model. The letters indicate mean flight height categories that were not significantly different in a pairwise comparison of means test based on the 95% credible interval around each modelled mean flight height. (b). Mean modelled flight height across the night.

### Flights at higher altitudes

The maximum modelled flight height estimate was 92.7 m with a maximum 95% CI of 144.1 m. Southern Bent-wing Bats took short flights at altitudes above 70 m over multiple minutes (observed for 5 - 7 minutes) during the night (Figure 3). This behaviour was observed on two instances from two different bats. These forays into higher airspace were observed from the 11 occasions (from 9 bats) when one-minute interval data was collected in the late summer period from the Portland subpopulation. One minute sampling periods only lasted for between 24 and 241 minutes due to the limited capability of the GPS trackers. This higher frequency sampling also showed that bats are at times regularly changing flight altitude up to a height of almost 40 m when not undertaking these higher-level flights.

**Figure 3.**
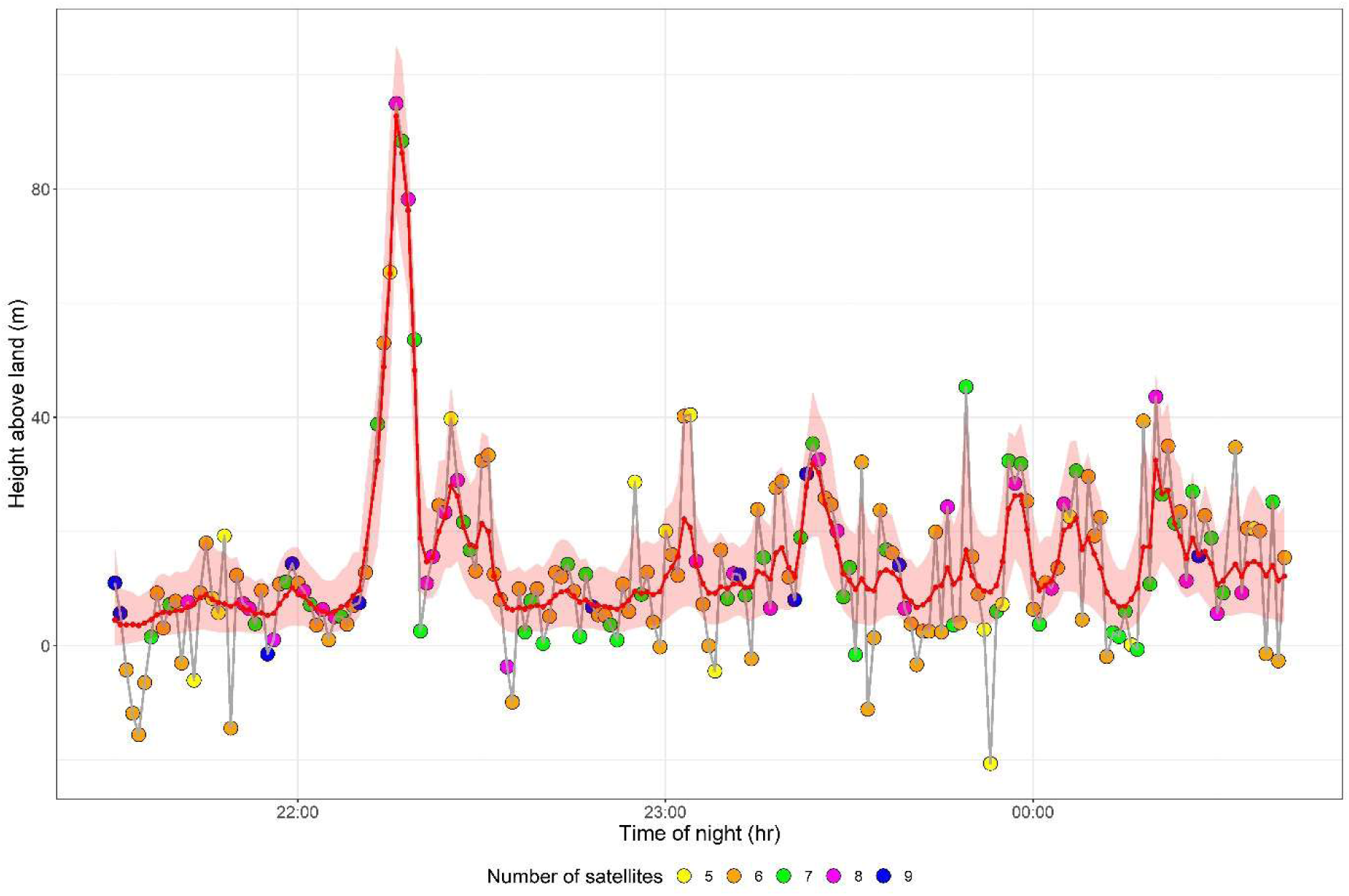
An example of raw and modelled flight heights above the ground surface illustrating a short higher altitude flight and frequent lower altitude changes in height. Raw bat flight height points are coloured by the number of satellites detected with the associated modelled heights in red. The shading shows the range of the 95% credible intervals for each modelled point.

### Correlation between successive flight heights

The model estimated significant autocorrelation in the AR1 flight height model, which declined with the time interval between successive observations. The autoregressive parameter ranged from approximately 0.78 for observations 1-minute apart to approximately 0.48 for observations 60-minutes apart, and 0.02 for observations 480 minutes apart (Figure 4Figure 4).

**Figure 4.**
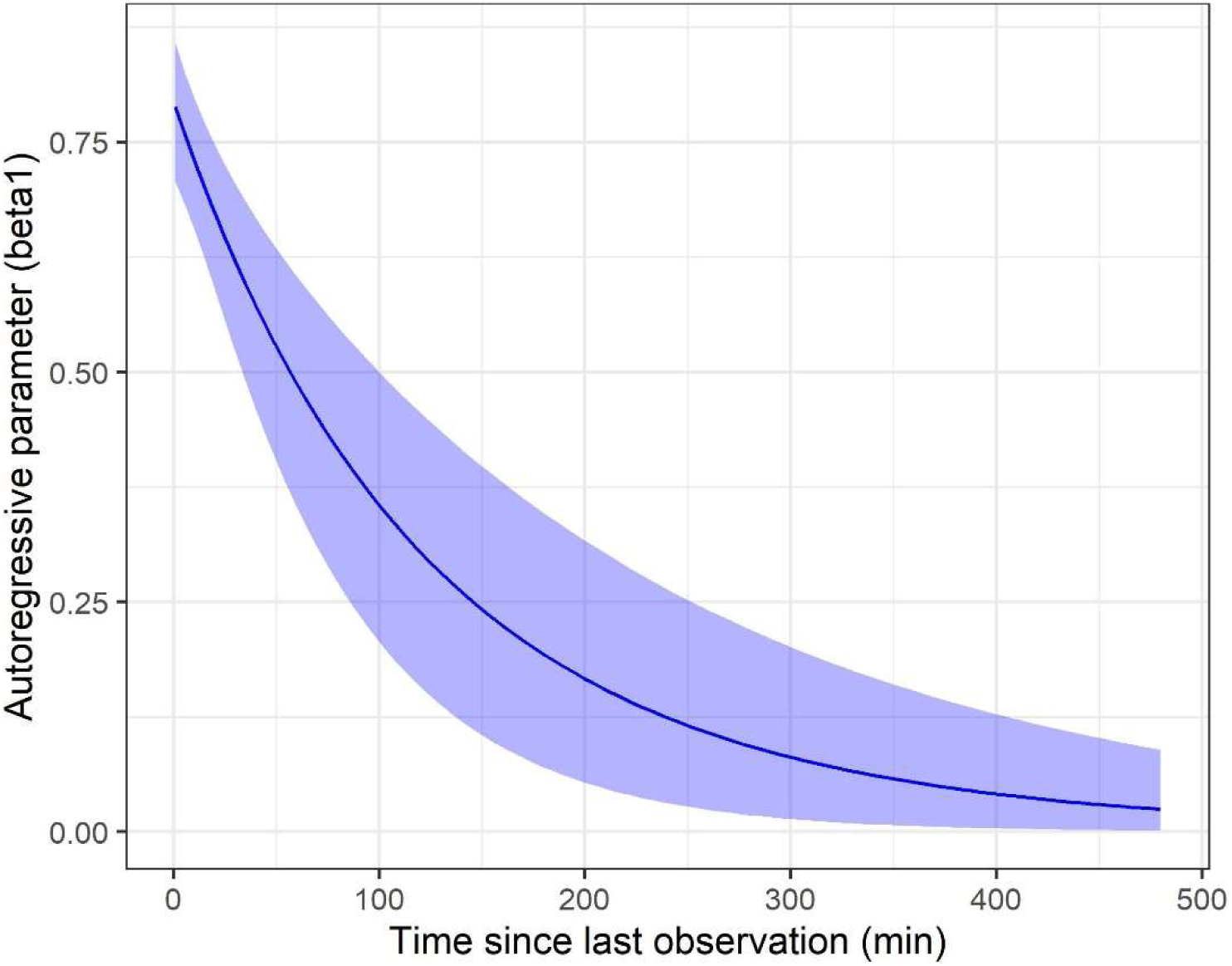
The autoregressive parameter generated from the AR1 model showing the high correlation between all GPS tracking time intervals in this study (i.e. 1-minute to 480 minutes).

## DISCUSSION

Understanding the flight height distributions for small bats is difficult because their small size limits the capability of suitable measurement devices. Yet, it is important to know what heights they fly at in order to help inform wind turbine collision risk. In this study we have demonstrated that small GPS units can provide an estimate of flight height patterns when modelled using a Bayesian state-space model. Our model showed that, in late summer and early spring, SBWB were most active below 30 m, with a general trend of a reduction in activity with increasing height, although some individuals were recorded flying more than 80 m above the ground. The reduction in activity with increasing flight heights identified here is similar to what has been reported for other small (<20 g) insectivorous bats, including other *Miniopterus* species based on acoustic studies ^16,57^.

In summer, modelled mean flight heights were higher in treed areas compared to untreed areas.

### Bat flight heights

GPS units used for wildlife tracking provide altitude measurements, which are influenced by actual flight heights of the individuals and the unit’s recording accuracy. Our flight height ranges were comparable to those recorded for other *Miniopterus* species. For example, calls from the Eastern Bent-wing Bat (*Miniopterus orianae oceanensis*) have been identified from acoustic recorder microphones set at 100 m above ground on a tethered static balloon, and Schreiber’s Long-fingered Bat (*Miniopterus schreibersii*) has been recorded between 4 and 85 m above ground on acoustic recorder microphones attached to static structures^16,57^. However, data collection for both of these studies was restricted to open and semi-open areas, and we were able to record bats at heights greater than those estimated from the acoustic recordings alone (i.e. a maximum modelled estimate of 92.7 m with a maximum 95% CI of 144.1 m). GPS units are also not subject to the well-documented limitations of acoustic recorders being deployed at height, which can significantly reduce the proportion of identifiable calls compared to ground-level recordings ^58^. It is well recognised that the number of individuals using an area cannot be determined from acoustic recordings ^59^, whereas our study had a known sample of individuals.

Insectivorous bats use flight for commuting, but the majority of their time in flight is spent foraging ^60,61^. This flexibility of SBWBs to use different height strata may provide access to food resources that would otherwise be unavailable. Prey type and abundance likely influence foraging habitat selection ^14^, with some insectivorous bats moderating their flight height based on the presence and availability of prey species. Species such as the Mexican Free-tailed Bat (*Tadarida brasiliensis*) feed on migrating insects at heights between 400 to 500 m during periods of nocturnal insect migration^62^. Other bat species make short excursions into higher airspace to forage to above 300 m in elevation ^31^. During summer and early autumn, the SBWB feeds primarily on moth species, whose larval stages are associated with both agriculture and native vegetation ^63^, however there is no available information on the flight heights of these moths in their adult form. We collected data for three weeks in February and March, and a similar period in September and October. If SBWBs forage on a different suite of insects, that fly at different heights at other times of the year or during wetter or drier periods, then their flight height patterns may vary across time.

### SBWB flight height and habitat

We found that modelled mean flight heights were greater when bat fixes were associated with treed compared to untreed habitat (13.8 m versus 7.3m for males and 11.4 m versus 7.1 m respectively for females), in summer. However, the range of flight height estimates from the modelled height data was broadly similar (2.00 m to 76.21 m for treed and 1.07 m – 92.7 m for untreed habitat). The degree to which bats forage above or below forest canopies, and whether or not this varies by vegetation type and structure, is poorly understood ^14^ and largely unknown for the SBWB. In this area mature trees are likely to be taller than 20 m but the treed classification also includes shrubs that are likely to be shorter. As such the range of modelled heights suggest that some flights are likely to be above the canopy, and some below or potentially adjacent to vegetation. Bats may have access to different resources above or below the canopy but may also have different constraints like difficulties with flying in cluttered areas. They may also at times fly along the edges of vegetation as has been observed with other species. The resolution (30 m) of the available vegetation data used during classification makes it difficult to know if bats are flying within or adjacent to vegetation if points are nearby, and this resolution also impacts the accuracy of division of flight heights between treed and untreed habitats, particularly in more patchy areas of the landscape.

We included sex as a variable in our analysis as males and breeding females will have differing energy requirements in the two seasons of our study, possibly leading to some level of variation in foraging strategy. SBWB undergo delayed implantation and breeding females would have been in the early stages of pregnancy in September/October ^64^ after reduced activity levels over winter ^42^. In February/ March the females had recently finished lactating and were in the process of increasing their body weight prior to winter. There does not appear to be much difference between the sexes overall, so although they may have different energy requirements this does not seem to be influencing their flight heights. One potential confounding factor is that first year females do not appear to breed and therefore may have energy requirements more similar to males^65^. When selecting individuals for tracking in the late summer period it was not always possible to determine if the females had bred that season, so female data were pooled. There have been few studies examining the flight height differences between the sexes for small insectivorous bats due to the very recent development of suitable technologies. Female Common Noctules (*Nyctalus noctula*), a larger open-space foraging bat (17 - 44 g)^66^, were found to fly higher than males above open areas in summer, although this may have been confounded by the data being collected in disparate time periods ^30^.

We found some difference in the mean flight heights of bats between spring and summer. For males, mean estimated flight heights in untreed areas were greater in summer than spring. Seasonal changes in flight distances and foraging effort have been observed for other species of insectivorous bats ^61^ but studies focused on seasonal changes in flight height of small insectivorous bats are lacking. Internationally, rates of reported bat mortalities from wind turbine collision are greater in late summer and autumn ^67^, as is the case in Victoria, including for the SBWB (unpublished data). This has at times been linked to a greater attraction of some species to turbines ^68^ at particular times of the year, however, if bats were flying higher during the late summer and autumn period this could also contribute to this pattern.

### Factors influencing modelled results

Our model assumes that all vertical error is attributable to the GPS but in fact there are other factors including the DEM error that are not considered. Furthermore, vertical error could be generated from poor horizontal accuracy of datapoints, causing the corresponding ground elevation estimate to be extracted at an incorrect location. We accounted for significant horizontal error by discarding all data with a HDOP greater than six.

There was a high correlation between consecutive modelled points derived from the AR1 model at shorter time intervals, with a gradual decline with increasing time interval. This may be due to a true highly associated flight pattern over longer periods resulting from factors like wind speed or direction. The high correlation may also be derived from factors unrelated to bat flight height like the orientation of satellites relative to each other during the controlled drone trials. It is also uncertain to what extent such a high correlation between consecutive samples is applicable to the bat data. SBWBs can fly more than 35 km in 60 minutes (unpublished data) in a very patchy landscape, so such a strong correlation is surprising, especially given the findings relating to differences between mean flight heights over trees and untreed areas in summer, and the frequent height changes observed during the one-minute sampling intervals.

The drone testing revealed that the number of satellites was a good predictor of vertical height error, at least in this controlled test environment. Measures of DOP (both horizontal and vertical when available) incorporate both the number of satellites and their position relative to the GPS unit ^35^, with configuration being important for determining vertical height accuracy. Some larger GPS units provide a VDOP error estimate that would better indicate the configuration of detected satellites than HDOP alone ^34^ and may have been a better predictor of vertical error however this is not a feature of the Lotek Pinpoint 10 GPS units.

It is not possible to truly emulate the full suite of conditions that would be contributing to measurement error from the free-flying bats in a controlled drone trial. Sixty-two percent of bat data locations were associated with trees, yet our drone flights all occurred in the open for safety reasons. Therefore, the relationship between vertical error, proximity to trees and number of detected satellites could not be established and is not considered in the model. The level of canopy closure is known to affect GPS accuracy by obscuring contact with satellites largely based on number of satellites detected ^32^. Our trials all occurred across relatively flat terrain in two confined 500 x 500 m areas, across up to 5.75 hours in a three-day period. It takes a complete satellite orbit of 11 hours and 50 minutes to experience all different satellite orientations relative to each other at a particular point on the land ^69^, so we tested our GPS trackers for only about half of a complete satellite orbit.

Our bat sampling frequencies were variable, ranging from 1-minute to 60-minute sampling intervals, with 54.6% being from 60-minute intervals. Shorter GPS sampling intervals (i.e. 2-3 s) are thought to yield the least vertical error ^28^, at least in stationary trials at ground level. Drone trials with variable sampling intervals (1-minute, 2-minute, 7-minute and 15 minute) showed no significant difference in vertical measurement error but we were unable to test longer sampling intervals in our drone trials to see if they contributed higher vertical uncertainty.

## CONCLUSIONS

Southern Bent-wing Bats are capable of flying to heights of more than 70 m and potentially up to 144 m (based on the upper bounds of the model estimates) above the ground at times, and can change height from near ground level to around 40 m within minutes. These flight patterns have implications for this species, given the complete overlap between renewable energy zones and their limited geographical range. There are currently more than 600 operational turbines within their range with hundreds more in the planning stage, and our results (in addition to existing mortality data^45^) indicate that the bat’s flight patterns are likely to be exposing them to collision risk with operational turbines at times. Therefore, to reduce the impacts on this Critically Endangered species, especially within the context of ‘no net loss’ or ‘nature positive’ settings, a range of mitigation measures are likely to be required, especially ones that have been shown to be effective for reducing bat mortalities, such as low wind speed turbine curtailment ^70^ .

## Ethics approval

Research was undertaken under ethics approvals S-2020-094 and S-2022-078 granted by the University of Adelaide, Animal Ethics Committee for the Warrnambool and Naracoorte subpopulations.

Research was undertaken with the Portland subpopulation under 23-005 assessed and approved by the Department of Environment, Energy and Climate Change, Animal Ethics Committee and Research Permit number 10010828.

## Competing Interests

### The authors declare they have no competing interests. <u>Funding</u>

- Department of Energy, Environment and Climate Action provided funding for drone testing of GPS units and assistance with manuscript preparation.
- Bat research was supported by the Victorian Speleological Society grant (Warrnambool subpopulation), Department of Environment, Land, Water and Planning Barwon South West Regions (Warrnambool subpopulation), Friends of Naracoorte Caves, Friends of Parks and Nature grant (Naracoorte subpopulation), Nature Fund, the Saving Native Species Fund and Southwest Environmental Alliance Landcare group, through the Glenelg Hopkins CMA and Department of Climate Change, Energy, the Environment and Water project funding (Portland subpopulation) and University of Adelaide student funds.
- The primary author received student funds assistance from the University of Adelaide.

### Authors’ contributions

AB led the project, sourced grant funding, collected bat and drone data, supported the analysis and wrote most of the manuscript. TP and LL provided guidance in project design. TP conducted project analysis with support from AB and contributed to manuscript writing. LL assisted with data collection and grant funding. All authors contributed to manuscript revision and approved final manuscript.

## Acknowledgements

The authors wish to acknowledge the generous contributions of the many volunteers who assisted with catching bats, attaching trackers and tracker retrieval for download. We would particularly like to acknowledge the assistance provided by Reto Zollinger, Yvonne Ingeme, Diane Gillatt, Dennis Matthews, Nicola Bail and Cassie Hlava. An earlier draft of the manuscript benefitted from review by Dr Andrew Bennett and Dr Pia Lentini.

Uncrewed Research Aircraft Facility at the University of Adelaide who organised trial sites and obtained all permissions and permits. Designed and constructed the custom-built mounting plate.

Pilot Dillon Campbell who flew the done for each trial and provided position output for inclusion into data analysis. Molly Ellis Hennekam, Steven Andriolo and Marnie Denlay for assistance with organisation and logistics of trials.

## List of abbreviations

DEMs: Digital elevation models
SBWB: Southern Bent-wing Bat

## Supplementary Material

**Figure S1.**
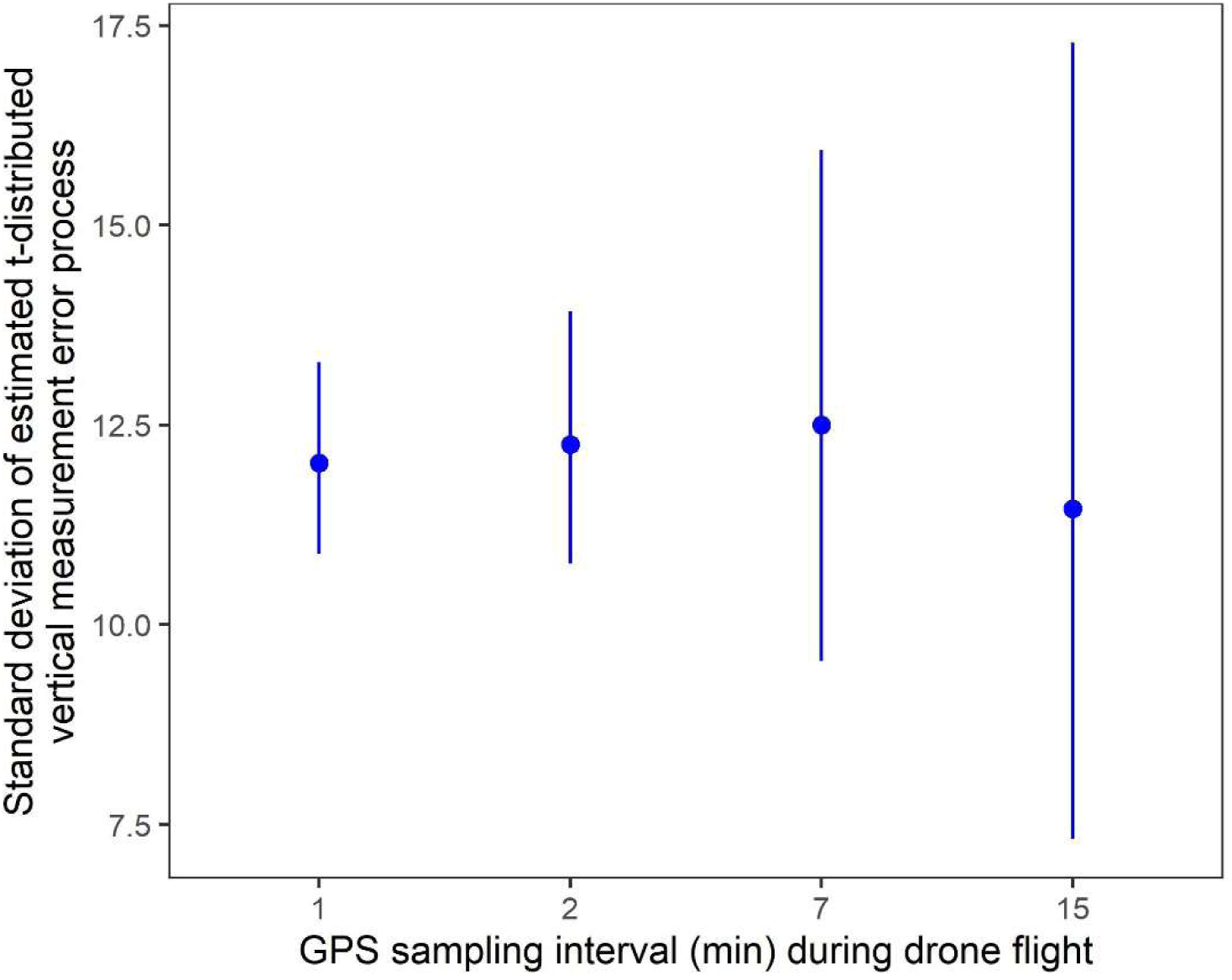
Comparison of the standard deviation of the vertical measurement error process for Lotek GPS units operating at different sampling intervals, estimated from data obtained by the drone trials. These results are conditional on a mid-range value of 6 satellites contacted for a location fix.

